# Protein allocation and enzymatic constraints explain *Escherichia coli* wildtype and mutant phenotypes

**DOI:** 10.1101/2020.02.10.941294

**Authors:** Tobias B. Alter, Lars M. Blank, Birgitta E. Ebert

**Author notes:** **Correspondence**: Tobias B. Alter, Institute of Applied Microbiology (iAMB), Aachen Biology and Biotechnology (ABBt), RWTH Aachen University, 52065 Aachen, Germany. **Funding information**: German federal and state governments (Grant-ID: PFLS015); CSIRO-UQ Synthetic Biology Alliance; German Research Foundation (DFG), Clusters of Excellence 236 “TMFB” and 2186„ The Fuel Science Center”.

## Abstract

Proteins have generally been recognized to constitute the key cellular component in shaping microbial phenotypes. Due to limited cellular resources and space, optimal allocation of proteins is crucial for microbes to facilitate maximum proliferation rates while allowing a flexible response to environmental changes. Regulatory patterns of protein allocation were utilized to account for the condition-dependent proteome in a genome-scale metabolic reconstruction of *Escherichia coli* by linearly linking mass concentrations of protein sectors and single metabolic enzymes to flux variables. The resulting protein allocation model (PAM) correctly approximates wildtype phenotypes and flux distributions for various substrates, even under data scarcity. Moreover, we showed the ability of the PAM to predict metabolic responses of single gene deletion mutants by additionally assuming growth-limiting, transcriptional restrictions. Thus, we promote the integration of protein allocation constraints into classical constraint-based models to foster their predictive capabilities and application for strain analysis and metabolic engineering purposes.

## 1 INTRODUCTION

For many decades, metabolic models have been developed to describe, unravel, and understand the drivers of microbial phenotypic behavior. In their simplest forms, these models quantitatively connect extracellularly observable phenomena leading to seminal empirical growth laws such as the hyperbolic relation between microbial growth and the substrate concentration, which is captured by the Monod equation (Monod, 1949). In general, coarse-grained models aided in explaining the dependencies between intracellular processes and corresponding phenotypes (Klumpp et al., 2009; Scott et al., 2010; You et al., 2013; Scott et al., 2014; Weiße et al., 2015; Erickson et al., 2017). Constraint-based modeling techniques, namely flux balance analysis (Savinell and Palsson, 1992), facilitate the prediction of growth rates and by-product secretion, as well as the investigation of metabolic flux distributions solely based on the stoichiometry of biochemical reaction networks and an appropriate cellular objective function (Varma et al., 1993; Varma and Palsson, 1994; Edwards and Palsson, 2000; Schuetz et al., 2007). The accompanied development of genome-scale constraint-based models (GEM) fostered the investigation of fundamental biological phenomena (Varma et al., 1993; Pramanik and Keasling, 1998; Mahadevan et al., 2002), the systematic analysis of complex omics data sets (Palsson, 2002; Blank et al., 2005; Becker and Palsson, 2008; Lewis et al., 2010) or the suggestion of favorable genetic perturbations for the overproduction of desired chemicals (Burgard and Maranas, 2003; Rocha et al., 2010; Cardoso et al., 2018; Alter et al., 2018; Alter and Ebert, 2019).

While the utilization of GEMs faciliated valuable contributions to hypothesis generation, explanation of physiological phenomena or optimal metabolic designs, the GEM’s predictive capabilities of microbial phenotypes strongly rely on *ad hoc* capacity bounds on key reactions (Palsson, 2002; Covert et al., 2003), without which basic phenomena, such as overflow metabolism under rich conditions are not observable *in silico*. The consideration of additional cellular processes and properties in metabolic reconstructions resolved these predictive insufficiencies of GEMs. In this manner, macromolecular expression (ME) models couple metabolism to gene expression by linking enzyme concentrations to metabolic reactions as well as accounting for the transcriptional and translational processes leading to the expression of enzymes (Lerman et al., 2012). For various steady-state environments, ME models simultaneously simulate maximum growth and substrate uptake rates, the underlying responses on the mRNA level as well as the corresponding gene expression profiles (O’Brien et al., 2013). Thus, ME models facilitate holistic insights into intracellular processes and how they are affected by environmental, biochemical, or genetic perturbations (Yang et al., 2016, 2019; Chen and Nielsen, 2019), while reliably informing about corresponding flux distributions. However advantageous ME models are for making correct predictions on a flux or phenotypic level, the detail and complexity of the represented network may be cumbersome regarding future applications in strain design approaches.

Another approach focuses on introducing enzyme limitations on fluxes by basic thermodynamic constraints. For *Saccharomyces cerevisiae* and *Escherichia coli*, the fusion of a thermodynamic and metabolic network model including an upper bound on total cellular Gibbs energy dissipation proved to explain metabolic realizations at various growth states, including maximum growth (Niebel et al., 2019). At the maximum rate of total Gibbs energy dissipation, the cell reorganizes its flux distribution, hence activates fermentation pathways to facilitate optimal growth with increasing substrate uptake rates at the expense of biomass yield. Classically, this observed trade-off between biomass yield and growth rate is explained on a structural rather than a thermodynamic level. For example, by assuming an upper limit for the intracellular concentration of enzymes, the limiting effect of cytoplasmic molecular crowding explained observed maximum growth rates and substrate uptake hierarchies (Beg et al., 2007; Vazquez et al., 2008). A similar principle emerged from the investigation of how the finite protein capacity of cellular membranes affects metabolism (Szenk et al., 2017). Here, a coarse-grained model derived from the physical description of protein repulsion on membranes correctly predicted the onset of overflow metabolism at the crowding limit of respiratory transmembrane proteins. In contrast, fine-grained models of unicellular organisms, such as resource allocation (Goelzer and Fromion, 2011) and whole-cell (Karr et al., 2012) models as well as self-replicating systems (Molenaar et al., 2009), uncover the traits and limits of metabolism by describing the complex management of resources among cellular subsystems. Despite disclosing the cellular economic principles that govern observed growth behavior, diverse applicability of fine-grained biological models, for example in biological engineering or data analysis disciplines, is hampered by the need for a tremendous number of mostly unknown parameters.

Cellular protein allocation and its regulation have previously been suggested as the main drivers of metabolic phenomena and a key process behind bacterial growth laws (Scott et al., 2010, 2014; Weiße et al., 2015; Mori et al., 2016; Noor et al., 2016; Erickson et al., 2017; Mori et al., 2019). The incorporation of protein constraints in GEMs exploits the principles of protein allocation as a fundamental growth law and simultaneously allows for the use of established, tractable, and intuitive constraint-based modeling methods. In this regard, by dividing the limited proteome into three growth-variant sectors representing (1) ribosomal proteins, (2) biosynthetic enzymes, (3) catabolically active enzymes, and one invariant housekeeping protein sector, the constrained allocation flux balance analysis (CAFBA) framework computes the optimal partitioning between these proteome sectors and the correspondingly weighted flux rates to reach maximum growth (Mori et al., 2016, 2019). Thus, CAFBA accounts for the trade-off between a limited protein availability for biosynthesis and growth, and enables a quantitative prediction of pathway usages under various conditions, particularly of sub-optimal growth yields as a consequence of overflow metabolism. To facilitate the utilization and analysis of proteomics data, enzyme kinetics were integrated into a GEM of *S. cerevisiae* and *E. coli* in the form of explicit enzymatic constraints on flux rates (Sánchez et al., 2017). The respective GECKO framework (GEM with enzymatic constraints using kinetic and omics data) gives detailed insights into metabolic realizations based on proteome measurements and predicts growth phenomena even without augmented (proteomic) data.

Here, we introduce an approach that consolidates protein allocation and enzymatic constraints on metabolic fluxes of an *E. coli* GEM. The resulting model, to which we will refer as the protein allocation model (PAM), accounts for the condition-dependent proteome (Schmidt et al., 2016), where enzyme kinetics are explicitly parametrized in the GECKO fashion (Sánchez et al., 2017). Proteome sectors that are not associated with metabolic reactions in the basic GEM are modeled by simple linear relations deduced from comprehensive measurements of the *E. coli* proteome (Schmidt et al., 2016). Assuming a constant protein concentration by mass, the PAM predicts experimentally observed phenotypes and intracellular flux distributions at maximum growth as well as carbon-limited conditions on various media. Beside the wildtype behavior, the predictive capability of the PAM is also demonstrated for strains harboring gene deletions or expressing heterologous proteins, without the need for laborious parameter sampling or explicitly constraining model variables. Instead, the observed phenomena are traced back to restrictions in the expression of metabolically active enzymes, presumably due to strictly determined responses of the transcriptional regulatory network. In line with previous studies, we emphasize the fundamental role of protein allocation in steering microbial metabolism. Moreover, the PAM framework appears to be a promising tool for the rational design of microbial production strains and, due to its constraint-based nature, enables rapid dry-lab screening of design and cultivation strategies with improved reliability.

## 2 RESULTS

### 2.1 Accounting for the total proteome in genome-scale metabolic models

A key challenge for microbes is the distribution of limited proteins on their intracellular processes to facilitate maximum growth under a given environmental condition. While the ribosomal content for the production of proteins is regulated to efficiency under mildly nutrient-limited to unlimited conditions (Scott et al., 2014; Bosdriesz et al., 2015; Mori et al., 2017), the protein household for energy and biomass precursor production generally contains unused or underused enzymes (O’Brien et al., 2016) to allow for a flexible response to changes in the environmental conditions. The three major protein sectors cover the total condition-dependent proteome and represent (1) ribosomal or translational protein, (2) metabolically active enzymes, and (3) unused as well as underused enzymes (Scott et al., 2010). We will refer to the latter as the excess enzyme sector for ease of simplicity.

To quantitatively account for the total condition-dependent proteome, we modeled and added each relevant protein sector independently to the *E. coli* K-12 MG1655 GEM *i*ML1515 (Monk et al., 2017) (Fig 1, cf. Methods section for a detailed description). Based on in-depth proteomics measurements of various *E. coli* strains grown under different conditions (Schmidt et al., 2016), the sum of the modeled protein mass concentration φ_P,c_ reflected in the data set was found to be constant among the tested strains and conditions (Fig 2A). By integrating equation (1) into the stoichiometric matrix, we accounted for this central microbial proteome feature in the PAM and fixed *φ*_P,c_ to 0.26 g 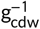, representing 81 % of the protein mass covered by the proteomics data set and 47 % of the approximated total cellular protein mass (Milo, 2013).

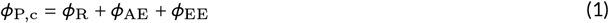

**FIGURE 1.**
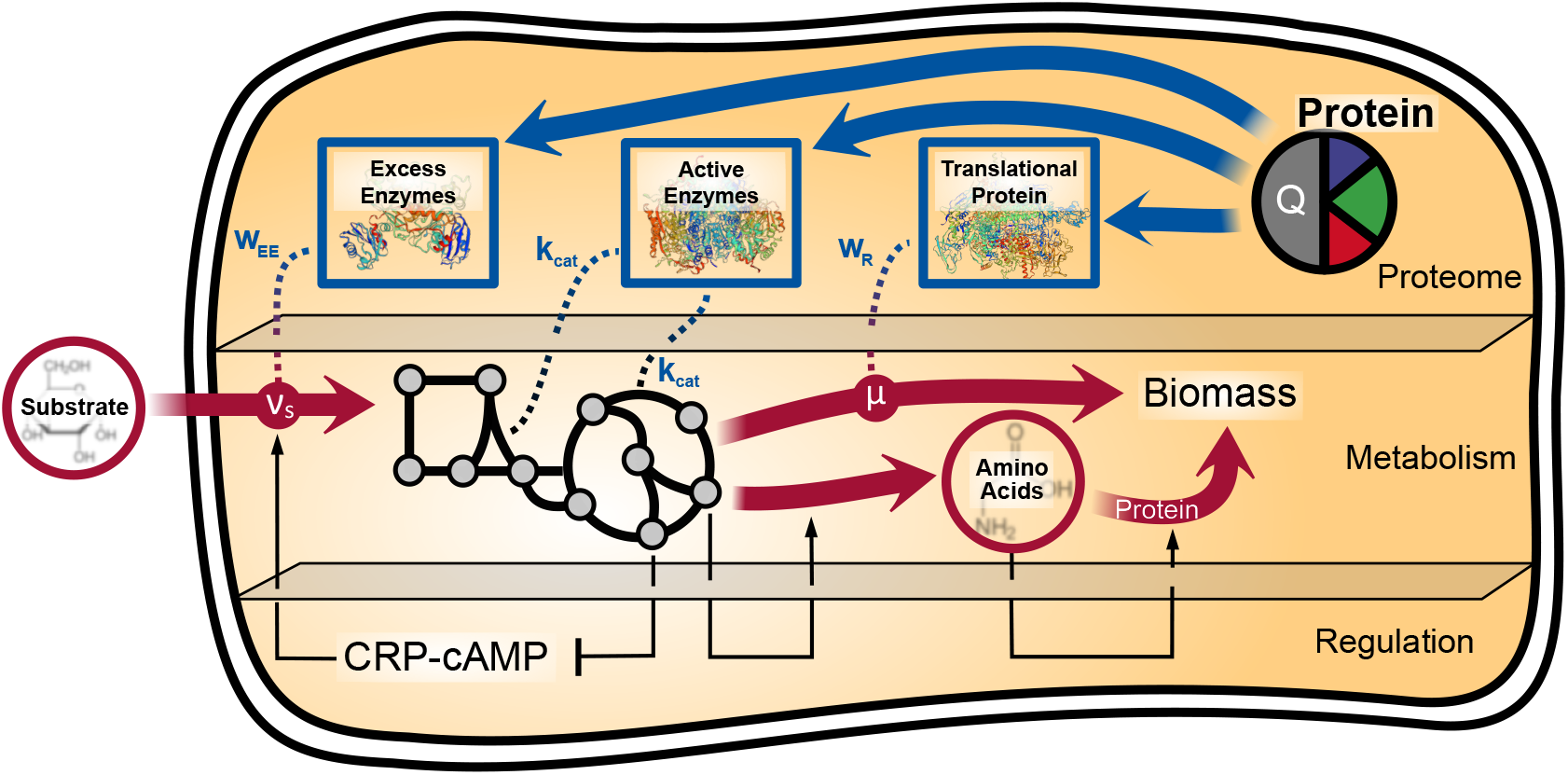
Scheme of the protein allocation model, including the classical metabolic as well as the added proteome level. *w*_R_, *w*_EE_, and *k*_cat_ represent the linear correlation factors between model variables and the translational, excess enzymes, and active enzymes sector, respectively. The dotted lines mark the model-inherent, linear relations between protein sectors and metabolic rates. A simplified mechanism of the regulation of microbial proteome, which is not part of the model, is additionally shown and was adapted from You et al. (2013). It depicts the activation of the expression of catabolically active enzymes by the CRP-cAMP complex, which synthesis is hampered by elevated amino acid precursor levels. Additionally, the activating effects of these precursors and amino acid concentrations on amino acid synthesis as well as protein synthesis, respectively, are sketched.

**FIGURE 2.**
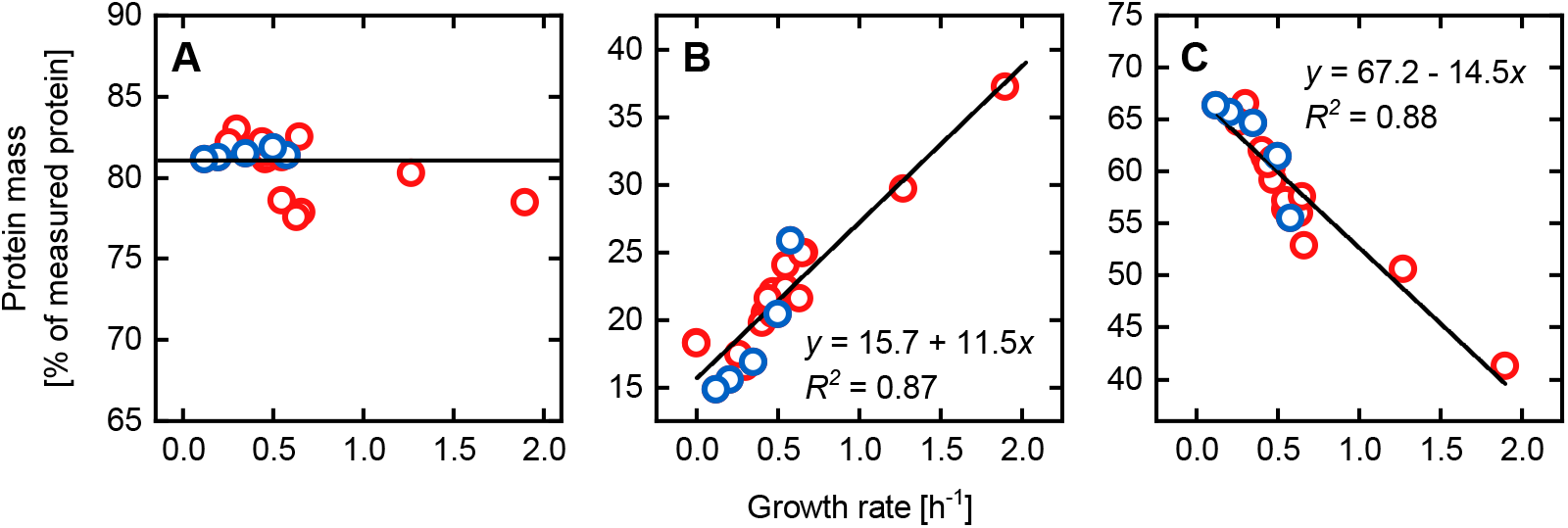
Experimentally determined protein masses of distinct proteome sectors (A-C) under several conditions. (A) Combination of the protein sectors shown in panel (B) and (C). The straight black line marks the 81 % of measured protein mass used to constrain protein availability in the PAM. (B) Translational or ribosomal sector including proteins assigned to the COG (Clusters of Orthologous Groups) class “translation, ribosomal structure and biogenesis” which are not covered by *i*ML1515. (C) All *i*ML1515 proteins found in the proteomics dataset. Black lines in (B) and (C) are linear fits of the data points resulting in the shown equations and coefficients of determination *R*^2^. Glucose chemostat and batch experiments are highlighted in blue and red, respectively. Data was taken from Schmidt et al. (2016).

Here, *φ*_R_, *φ*_AE_, and *φ*_EE_ are the protein mass concentrations for the ribosomal, the metabolically active enzyme, and the excess enzyme sectors, respectively.

#### 2.1.1 The ribosomal sector

In agreement with previous studies, a linear correlation between the ribosomal sector *φ*_R_ and the growth rate (Scott et al., 2014; Schmidt et al., 2016) was implemented and described as

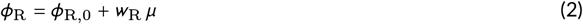

where the slope *w*_R_ compares with the inverse of the maximum ribosomal elongation rate and the intercept *φ*_R,0_ indicates an increasing overcapacity of ribosomes with a decreasing growth rate *μ* (Mori et al., 2017; Metzl-Raz et al., 2017). Both parameters *w*_R_ and *φ*_R,0_ were determined for *E. coli* by fitting equation (2) to measured, cross-conditional concentration data of the ribosomal proteome sector (Fig 2B). Consequently, *w*_R_ and *φ*_R,0_ were set to values of 50.0mg 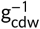 (9 % of the total protein mass) and 36.8 mg 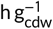, respectively.

#### 2.1.2 The excess enzyme sector

A major evolutionary characteristic of microbes facing unforeseeable changes in environmental conditions is the synthesis of excess proteins. This protein hedging empowers the cell to quickly ramp up central carbon metabolism fluxes upon a sudden increase in substrate availability or to immediately catabolize a new carbon source. However, such a stockpiling of proteins significantly reduces the growth rate (O’Brien et al., 2016). Protein expression and (over-)allocation is generally coordinated by the cyclic AMP (cAMP) dependent signaling pathway via the cAMP-activated global transcriptional receptor protein (CRP) (Kolb, 1993; You et al., 2013) (Fig 1). The CRP-cAMP complex enhances the transcription of over 100 genes by attaching near or at their promoter regions thereby mediating the binding of RNA polymerase for transcription initiation (Busby and Ebright, 1999). The target genes are mainly associated with catabolism, thus, CRP-cAMP aids in stimulating the carbon influx leading to the accumulation of precursors for amino acids, and therefore for protein synthesis. These precursors, such as oxaloacetate or α-ketoacids, inhibit the cAMP synthesis, thus lower the CRP-cAMP level which in turn closes the negative feedback loop between protein precursors and substrate uptake (we refer to You et al. (2013) for respective insights into the cAMP signaling pathway).In this way, CRP-cAMP indirectly coordinates the global protein allocation, including the allocation of excess enzymes, even though transcription of only a small fraction of genes is directly affected. Hence, by regarding the cAMP-controlled signaling pathway as a blueprint for protein synthesis regulation, we modeled the excess enzyme sector as a negative linear function of the substrate uptake flux, which we mathematically expressed as

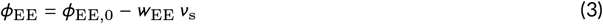

where φ_EE,0_ is the excess enzyme concentration at zero substrate uptake and *ν*_s_ the substrate uptake rate. *w*_EE_ relates the decrease of the excess enzymes concentration to an increase in *ν*_s_. *φ_EE,0_* was determined from a linear fit of proteomics data of all *i*ML1515 proteins, resulting in a value of 0.17g 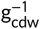 equivalent to 31 % of the total protein mass (Fig 2C).The slope w_EE_ is individually assigned for each substrate under the assumption that the excess enzyme concentration is zero at the maximum substrate uptake rate. Thus, *w*_EE_ is calculated from

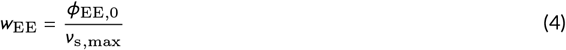

The maximum substrate uptake rate *ν*_s,max_ needs to be provided as an observable but can be directly inferred from the PAM by assuming an upper limit in the transcriptional capacity, as will be shown.

#### 2.1.3 The active enzyme sector

To account for the enzyme cost of metabolic fluxes, we integrated enzyme mass balances for all relevant metabolic reactions in the stoichiometric matrix of iML1515 according to the GECKO framework (Sánchez et al., 2017). The employed enzyme mass balance formulation (equation (5)) relates the flux *ν*_e_ to the minimally required concentration *ρ*_e_ of the enzyme catalyzing reaction e by the enzyme’s maximum turnover number *k*_cat,e_. Since the PAM accounts for the total proteome *by mass,* we transformed molar to mass concentrations using the molar mass *M*_e_ of each individual model enzyme (cf. the Methods section for a detailed description).

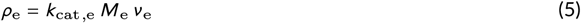

Eventually, the sum of the concentration of all *E* enzymes constitutes the metabolically active enzyme sector *φ*_AE_ and is expressed as

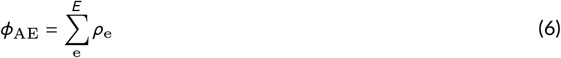

Parametrization of the active enzyme sector is of particular importance to facilitate a meaningful relation between fluxes and enzyme concentrations. For the PAM we determined *k*_cat_ values for 2843 protein-related reactions of iML1515 from queries of the databases BRENDA (Jeske et al., 2019), SABIO-RK (Wittig et al., 2012), and UniProt (Bateman, 2019) following the protocol of Sánchez et al. (2017). The *k*_cat_ dataset was manually curated to allow for the simulation of reasonable phenotypes (cf. additional files for the final dataset), thus transforming the primary *in vitro k*_cat_ estimates to effective (or apparent) *in vivo* turnover numbers *k*_app_ (Valgepea et al., 2013; Davidi et al., 2016). Recently, measurements of *k*_app,max_, the maximum *k*_app_ values across conditions, were extrapolated to genome-scale using machine learning models informed with biochemical and enzymatic data (Heckmann et al., 2018). Interestingly, although the distribution of *k*_app,max_ values is significantly different from our curated set (Fig 3), simulations using the PAM parametrized with either of both *k*_app,max_ sets yielded comparable phenotypes (data not shown). Thus, it appears that the ratio of *k*_cat_ values between different reactions or pathways is an important factor for the predictive capability of PAMs in general.

**FIGURE 3.**
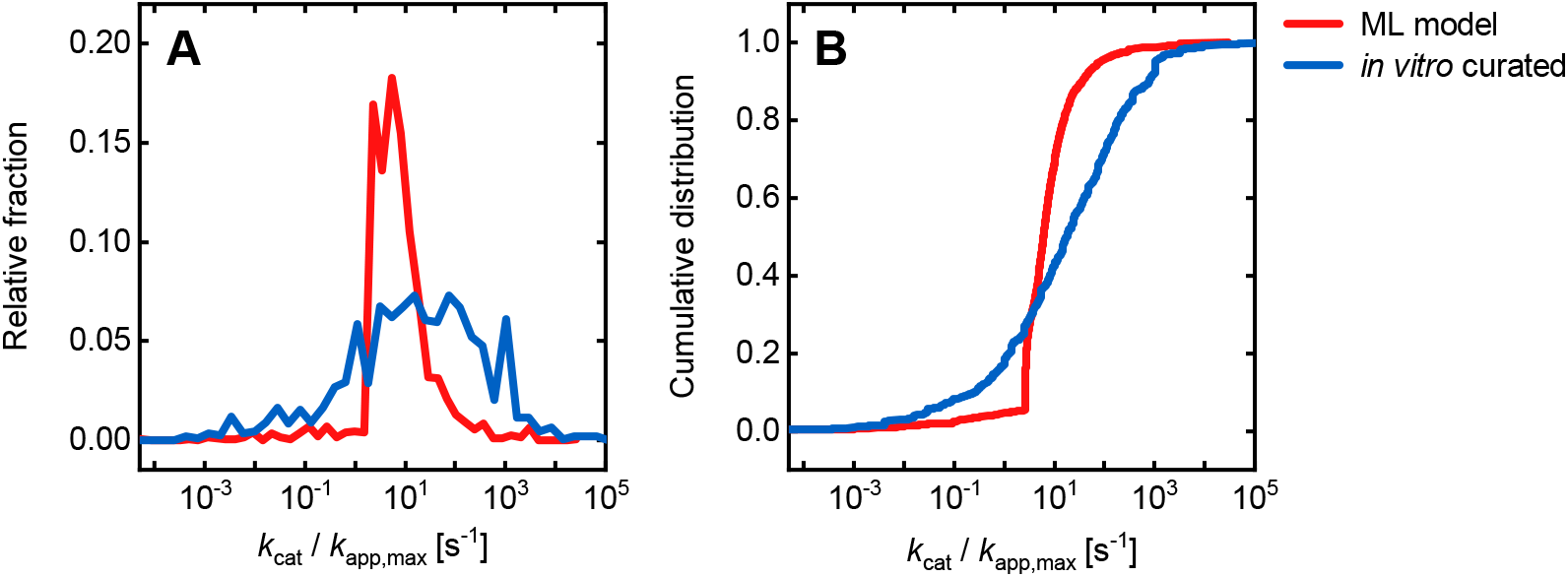
Relative fraction (A) and cumulative distribution (B) of *k*_cat_ *(in vitro* curated) and *k*_app,max_ (machine learning (ML) model) values. *k*_cat_ values are curated data extracted from the BRENDA, SABIO-RK, and UniProt databases, whereas k_app,max_ values were taken from Heckmann et al. (2018).

### 2.2 Protein allocation model predicts wildtype phenotypes and flux distributions

#### 2.2.1 Prediction of *E. coli* phenotypes

To benchmark the predictive capabilities of the PAM, wildtype phenotypic behavior on a glucose minimal medium was simulated and compared to extensive literature data (Perrenoud and Sauer, 2005; Nanchen et al., 2006; Vemuri et al., 2006; Valgepea et al., 2010; Folsom et al., 2014; McCloskey et al., 2014; Peebo et al., 2015; Folsom and Carlson, 2015). The maximum glucose uptake rate, a parameter for the excess enzyme sector, was set to 8.9 mmol 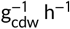 which supported a maximally observed growth rate of 0.65 h^-1^ (Perrenoud and Sauer, 2005). The simulated phenotypes are in good agreement with experimentally observed data for a range of carbon limited conditions and depict significant improvements compared to the purely stoichiometric model *i*ML1515 (Fig 4). Particularly the acetate secretion trend correctly mirrors the metabolic overflow characteristics of *E. coli* starting from glucose uptake rates above 4.3 mmol 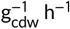 (Fig 4B). Despite reasonable projections of growth, acetate secretion, and oxygen uptake, the PAM as well as the *i*ML1515 model overestimate carbon dioxide secretion rates, pointing to potential inconsistencies in the carbon content of the model-inherent biomass equation.

**Figure 4.**
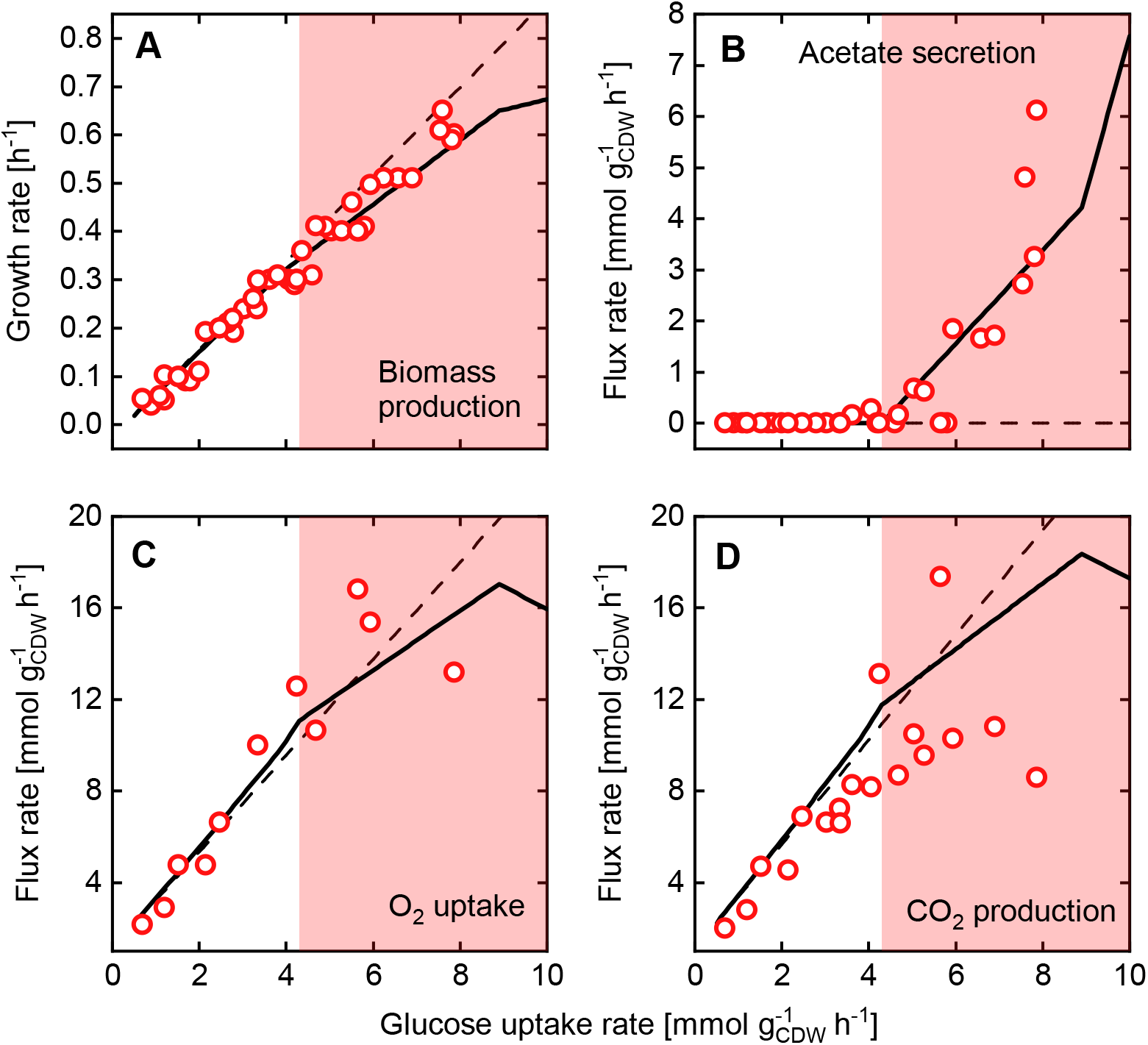
Phenotypic predictions of growth (A), acetate secretion (B), oxygen uptake (C), and carbon dioxide production (D) rates for *E. coli* for a range of physiologically relevant glucose uptake rates using the PAM (black lines) in comparison to data from the literature (red dots). Simulation results using the original stoichiometric *i*ML1515 model are additionally shown (dashed lines). The red shaded area marks the occurrence of fermentation resulting in the secretion of acetate.

The phenotypes simulated by the PAM are also mirrored in the fluxes through central metabolic pathways as shown in Fig 5. The distribution of carbon flux among the pathways qualitatively follows previous findings (Nanchen et al., 2006). Under carbon limited and fully respiratory conditions, flux rates through Embden-Meyerhof-Parnas (EMP) pathway tricarboxylic acid (TCA) cycle, pentose phosphate (PP) pathway, and the glyoxylate shunt (GLYXS) scale linearly with the glucose uptake rate. In the vicinity of the turning point from a fully respiratory to partially fermentative metabolism, the activity of the GLYXS diminishes completely. As soon as limitations in the global protein household impede exclusive ATP production via respiration, the carbon flux is partly diverted from the NADH-yielding, and thus respiration-fueling TCA cycle towards acetate. Simultaneously, the split between the EMP and PP pathway starts to increase with elevated glucose uptake rates in favor of glycolysis. Beyond the assumed maximum glucose uptake rate, the PAMs stoichiometric and protein allocation constraints still support feasible growth states. However, the increase in growth rate is bought at the expense of biomass yield, which becomes evident by a drastic decrease of the flux through the EMP pathway and the TCA cycle, as well as a progressively increasing acetate secretion rate.

**FIGURE 5.**
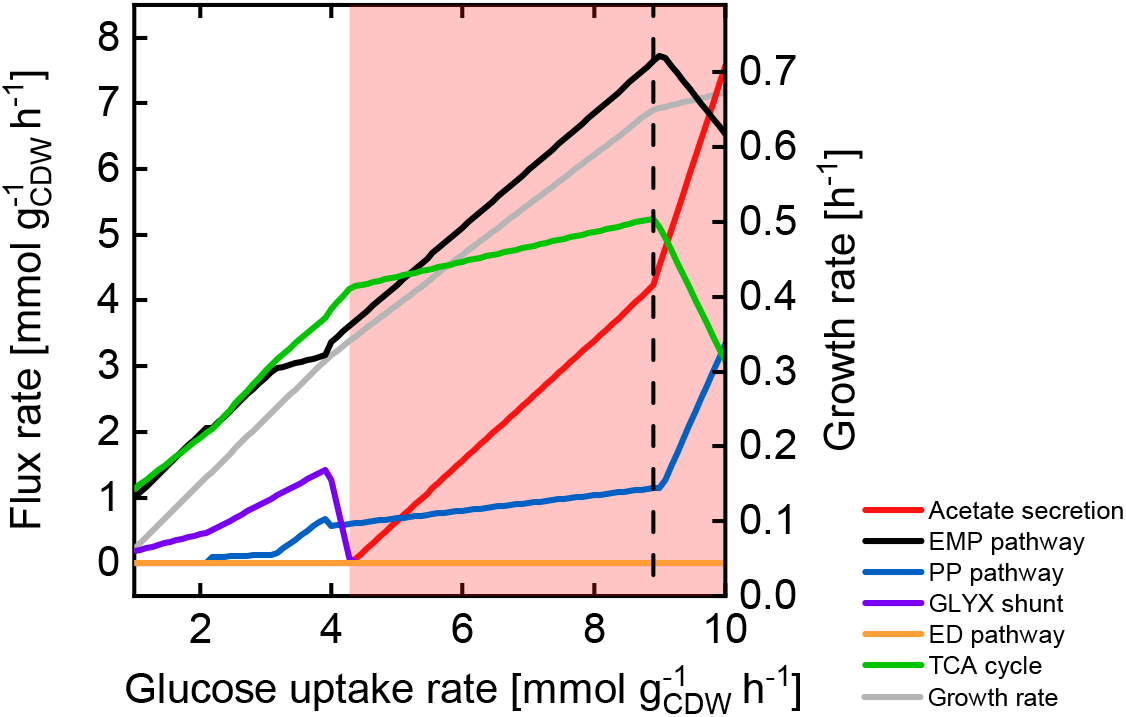
Simulated fluxes through central metabolic pathways for a range of glucose uptake rates by the PAM. The dashed line marks the maximum glucose uptake rate of 8.9 mmol 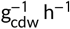 where excess enzyme concentration *φ*_EE_ becomes zero. The red shaded area marks the occurrence of fermentation resulting in the secretion of acetate.

This does not necessarily indicate an underestimated maximum glucose uptake rate since PAM parametrization stays feasible even for maximum glucose uptake rates far beyond physiologically relevant values (data not shown). In fact, this suggests that the metabolically limiting factor for *E. coli* is neither stoichiometry nor protein allocation. As we will show, maximal division rates may rather be attributed to a limited maximum number of proteins, thus pointing to transcriptional restrictions as the determining growth limiting factor.

#### 2.2.2 Computing maximum substrate uptake rates

Since glucose is *the* preferred carbon source of *E. coli,* its metabolism and regulation is adapted to effectively utilize glucose, which led us to assume that there are no excess enzymes under glucose-excess conditions. However, such a state of adaption does not hold for alternative carbon sources and thus, experimentally observed substrate uptake rates may not reflect growth states with (near-)zero excess enzymes. To enable the PAM to simulate growth on any alternative carbon source without the need for extensive cultivation data, we approximated maximum substrate uptake rates assuming a limited, total protein synthesis rate *N_P_*. *N_P_* represents the sum of molar synthesis rates of proteins from the active enzyme sector and the ribosomal sector and is calculated as

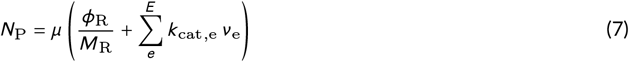

Here, *M*_R_ is the sum of molar masses of all 21 ribosomal subunits, which adds up to 3.5 × 10^5^ g mol^-1^. Based on the PAM and phenotypic data of *E. coli* grown on several non-glucose substrates (Gerosa et al., 2015), we found that a *N_P_* of 2.04 μmol 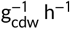 defines maximum substrate uptake rates (Table 1, cf. Methods section for a detailed description). Interestingly, the growth-optimal acetate uptake rate approaches a global maximum at 18.44 mmol 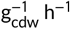 without reaching the upper bound for *N*_P_. Thus, other factors such as ATP generation must ultimately limit maximum growth on substrates, which, as in the case of acetate, prohibit the use of energy-yielding metabolic modes other than respiration. By parametrizing the excess enzyme sector (equation (3)) with the substrate-specific maximum substrate uptake rates, PAM simulations of phenotypes of *E. coli* grown on the individual alternative carbon sources showed a good correlation with experimentally observed data (Fig 6). The phenotypes predicted for the determined maximum substrate uptake rates generally overestimate the observed growth and acetate secretion rates (Appendix Fig S1), indicating an unused potential to adapt *E. coli* to these alternative carbon sources. Exemplarily, Fong et al. (2005) exploited the apparent potential for growth on glycerol by evolving the glycerol uptake rate to around 15 mmol 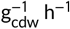, which is close to the PAM’s predictions.

**TABLE 1.** *In silico* determined maximum substrate uptake rates according to a maximally allowable total protein synthesis rate *N_P_* of 2.04 μmol 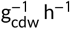. Since fully anaerobic conditions are infeasible for the PAM and also *i*ML1515, minimal oxygen uptake rates of 1.5 mmol 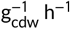 and 3.5 mmol 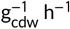 were used to simulate micro-aerobic growth on glucose and xylose, respectively.

| Substrate | Maximum uptake rate<br>[$\text{mmol g}_{\text{cdw}}^{-1} \text{h}^{-1}$] | Substrate | Maximum uptake rate<br>[$\text{mmol g}_{\text{cdw}}^{-1} \text{h}^{-1}$] |
| --- | --- | --- | --- |
| Glucose (aerobic) | 9.8 | Fructose | 9.6 |
| Glucose (micro-aerobic) | 20.3 | Fumarate | 26.5 |
| Galactose | 9.4 | Acetate | 18.4 |
| Glycerol | 14.9 | Gluconate | 9.5 |
| Pyruvate | 53.0 | Succinate | 24.0 |
| Xylose (aerobic) | 14.2 | Mannose | 6.4 |
| Xylose (micro-aerobic) | 26.6 | Glucosamine | 9.5 |

**FIGURE 6.**
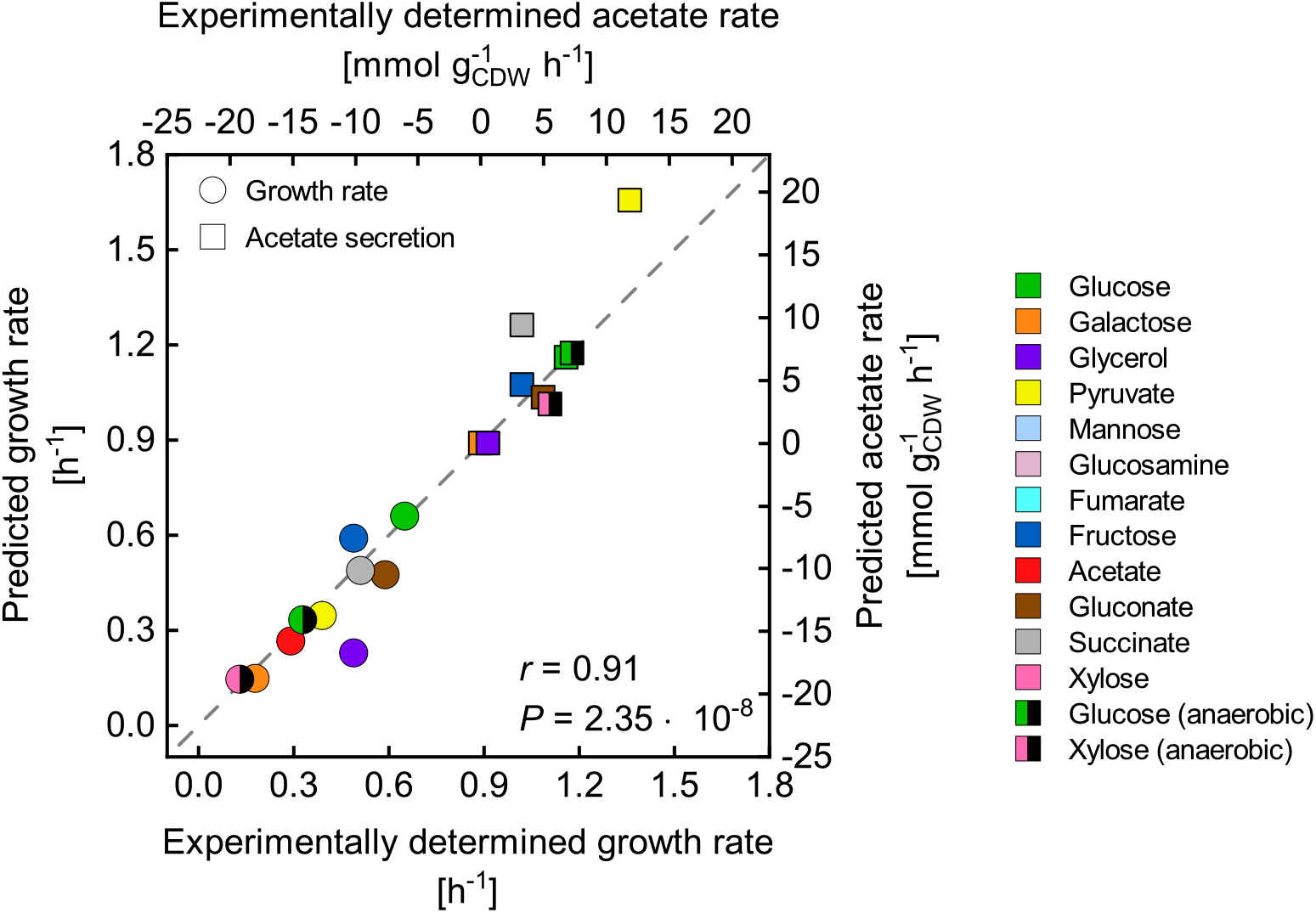
Comparison between experimentally determined (Gerosa et al., 2015) and predicted growth and acetate secretion rates using several substrates as sole carbon sources. Substrate uptake rates were constrained according to the reported values. Maximum substrate uptake rates were approximated according to a maximum total protein synthesis rate *N*_P_ (Table 1). The goodness of the correlation between simulations and experiments was determined using the Pearson correlation coefficient *r* and the corresponding *P* value.

#### 2.2.3 Prediction of *E. coli* flux distributions

By exploiting the optimality principles of microbial growth, GEMs give quantitative insights into the intracellular flux distribution and pathway usage based purely on stoichiometric constraints. For a particular environmental condition of interest, the prediction accuracy of the stoichiometric GEM generally scales with the amount and accuracy of experimental data introduced to the model in the form of flux constraints. With a minimum need for such data, the PAM allows for an accurate blueprint of the intracellular metabolic processes. A comparison with fluxomics data from multiple studies shows that flux distributions of the central carbon metabolism of *E. coli* grown on a minimal glucose medium are well predicted by the PAM, which is indicated by Pearson correlation coefficients up to 0.97 (Fig 7). Here, the PAM was constrained with the measured glucose uptake rates, and the excess enzyme sector was parameterized according to the methodically determined maximum glucose uptake rate, which, in turn, was a direct model output (cf. Section 2.2.2). Thus, protein allocation and enzymatic constraints alone enhance constraint-based modeling as a metabolic prediction tool, particularly under data scarcity. Yet, rather high discrepancies between simulated and experimental flux data from Nanchen et al. (2006) and Haverkorn van Rijsewijk et al. (2011) (Fig 7B and C) were found for the pyruvate kinase. These incongruities result from the neglect of the dephosphorylation of phosphoenolpyruvate (PEP) for the glucose uptake by the phosphotransferase system (PTS) by Nanchen et al. (2006), thus necessitating elevated pyruvate kinase fluxes to balance PEP.

**FIGURE 7.**
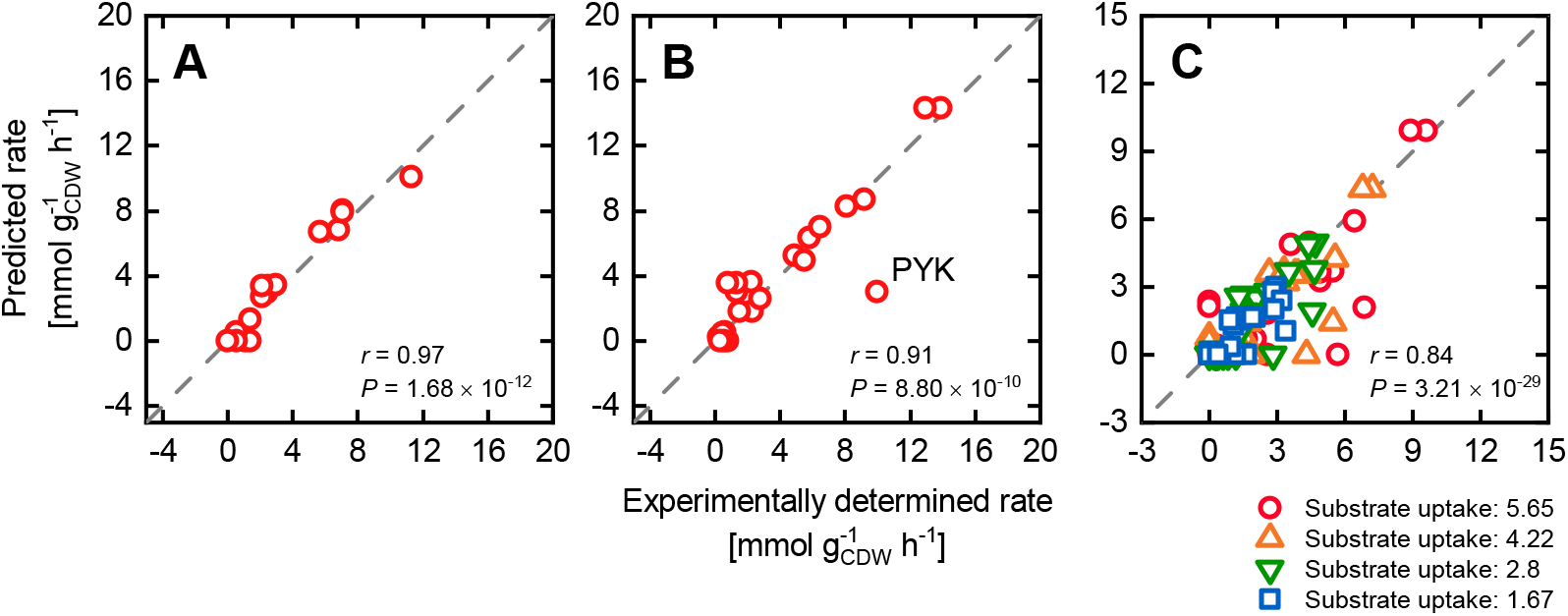
PAM predictions of intracellular fluxes of the central carbon metabolism of *E. coli* grown on a glucose minimal medium. The glucose uptake rate was constrained with experimentall determined values. The excess enzyme sector was parameterized according to the computationally determined maximum value of 9.82 mmol 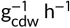. The predictions are compared with experimental flux data from Gerosa et al. (2015) (A), Haverkorn van Rijsewijk et al. (2011) (B), and Nanchen et al. (2006) (C). The goodness of the correlations was computed based on the Pearson correlation coefficient *r* and the corresponding *P* value.

Using the methodically determined maximum substrate uptake rates and constraining the PAM with measured uptake rates, flux distributions were approximated for non-glucose carbon sources (Fig 8). Here, the prediction capabilities were more diverse resulting in high correlations for acetate, galactose and succinate (*r* > 0.92) but intermediate to weak predictions, e.g., for fructose or gluconate (*r* < 0.65). In the case of gluconate consumption, the ED pathway was experimentally observed to be the main catabolic route, whereas the simulated carbon flux was exclusively channeled towards pyruvate through the pentose phosphate (PP) pathway. The flux split between the PP and ED pathway proves to be highly sensitive to the ratio in the protein demands of both pathways. Substrate-dependent differences in the biomass compositions as well as inconsistencies in the applied *k*_cat_ values, particularly of backward reactions, may cause the observed discrepancies between experimentally determined and simulated pathway usages.

**FIGURE 8.**
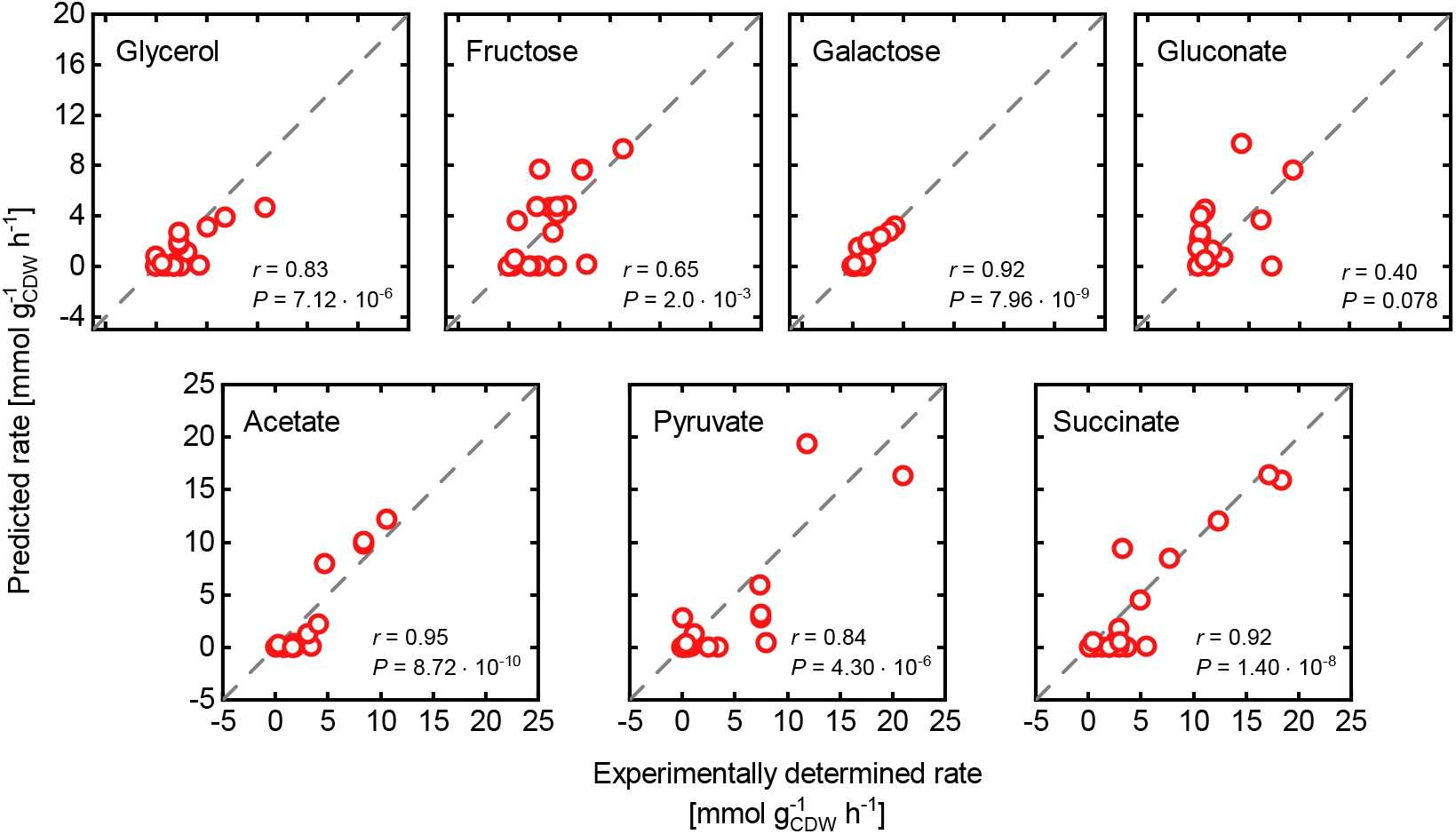
PAM predictions of intracellular fluxes of the central carbon metabolism of *E. coli* grown on several alternative carbon sources. The substrate uptake rates and the excess enzyme sector were constrained with the determined maximum values. The predictions are compared with experimental flux data from Gerosa et al. (2015). The goodness of the correlations was computed based on the Pearson correlation coefficient *r* and the corresponding *P* value.

### 2.3 PAM explains the growth defect upon heterologous protein expression

Enzyme overexpression or the expression of heterologous, non-native genes and their products is a common strategy in many biotechnological disciplines. General purposes are the introduction of novel cellular functionalities, flux enforcement through a specific pathway, or the investigation of cellular processes via reporter proteins. In any case, the introduced pull of proteins from the limited native protein household and the resulting metabolic burden inevitably causes a defect in growth.

To investigate and evaluate the response of the PAM to such an induced protein demand we simulated growth for a range of expression levels of an enhanced green fluorescent protein (eGFP) and compared relative growth rates with experimental data from Bienick et al. (2014). For the standard amount of total condition-dependent protein *φ*_P,c_ of 0.26 g 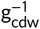 the simulated relative growth defect is significantly higher compared to experimentally observed growth for intracellular eGFP concentrations beyond 0.03g 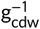 (Fig 9A). However, Bienick et al. (2014) employed an *E. coli*TUNER strain, a derivative of the genome-reduced BL21 strain optimized for protein expression. Thus, the protein demand for the growth rate-independent housekeeping sector is reduced and made available to metabolic, condition-dependent proteome sectors, which can be reflected in the PAM by an increase of *φ*_P,c_. As a result, by expanding *φ*_P, c_ about 20 %, the surplus in protein availability attenuates the detrimental effects of eGFP expression on growth and results in an excellent reproduction of experimentally observed phenotypes (Fig 9A) (cf. Methods for the determination of the optimal *φ*_P, c_).

**FIGURE 9.**
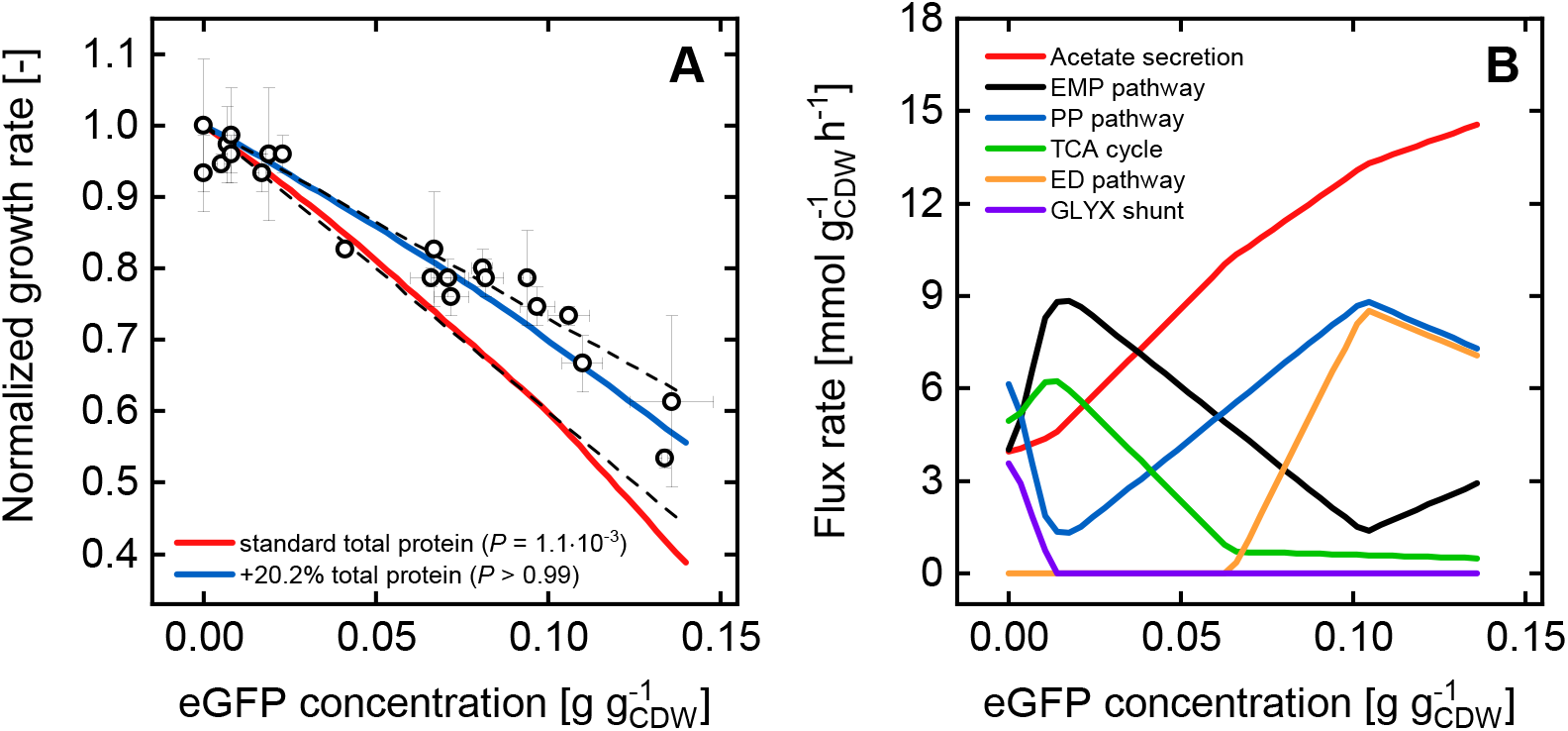
(A) Simulated growth rates relative to the maximum growth rate are shown for a range of intracellular eGFP concentrations using the PAM with a standard (red line) and increased (blue line) total condition dependent protein concentration *φ*_P, c_ (0.26g 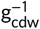 and 0.031g 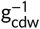 respectively). The shown experimental data is taken from Bienick et al. (2014). The given P values derived from a Student’s t-test, indicate how well the experimental data is explained by the model predictions. The dashed lines illustrate the predicted range of values based on theoretical considerations by Bienick et al. (2014). (B) Simulated fluxes through central metabolic pathways are shown for a concentration range of intracellularly expressed eGFP applying a *φ*_P,c_ of 0.31 g 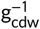.

Interestingly, the relation between eGFP expression strength and growth predicted by the PAM is non-linear, which is in contrast to a previous theoretical postulation (Bienick et al., 2014). The non-linearity arises from a combined effect of a protein drain from the ribosomal *and* the active metabolic enzyme sectors. An enforced protein waste causes a decline in the ribosome concentration resulting in a reduced translation rate. At the same time, protein allocation to the metabolic sectors producing biomass precursors, amino acids, and energy is limited, which, in a manner of a vicious cycle, further decelerates protein production and growth. This intracellular tug-of-war for proteins is intrinsically manifested in the PAM leading to an outperformance in the prediction of protein overexpression phenotypes over too simplified coarse-grained models. Moreover, the PAM discloses relevant effects of a protein drain on the pathway flux level (Fig 9B). For an increasing eGFP expression strength and an accompanied protein deficiency, central carbon fluxes are progressively diverted to fermentation pathways (acetate secretion) and eventually to the ED pathway. Both routes are more protein efficient in metabolizing carbon substrates but ultimately yield fewer energy equivalents per substrate molecule as compared to respiration or the EMP pathway, respectively (Flamholz et al., 2013; Mori et al., 2019; Ng et al., 2019).

### 2.4 Limitations in the protein allocation of single enzymes lead to gene deletion mutant phenotypes

Alongside the over- and heterologous expression of genes, rearrangement of metabolic networks and flux distributions by gene deletions is a core instrument in metabolic engineering. In recent years, the engineering of microbial cell factories has increasingly been supported by computational strain design methods which, besides elementary-mode analysis frameworks, utilize constraint-based models for describing the (mutant) metabolism. However, in contrast to the vast number of model-driven strain design and optimization methods Maia et al. (2016), constraint-based methods have often proven unreliable in predicting phenotypes of gene-deletion mutant strains (GMS). While GMSs have been shown to evolve towards FBA-predicted phenotypes Fong and Palsson (2004), observed growth defects and intracellular fluxes of non-evolved GMS can not be explained by stoichiometry and a cellular growth objective alone Kim and Reed (2012); Long et al. (2016); Long and Antoniewicz (2019a).

Firstly, we tested the impact of ten single-gene deletions on the PAM’s FBA results by parametrizing the excess enzyme sector with the methodically determined maximum glucose uptake rate of 9.82 mmol 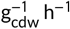. The calculated phenotypes did not significantly differ from the unperturbed wildtype solutions and hence, did not compare to experimental data from Fong et al. (2006) and Long et al. (2016) (Appendix Fig S2). Only by constraining the glucose uptake rate to the measured values, the simulated growth and acetate secretion rates correlate well with the experiments (Fig 10). Furthermore, a blind test, more precisely FBA simulations with augmented *in vivo* glucose uptake rates of the GMS but with an *intact* target gene, yielded the same phenotypes (Appendix Fig S3). These results led us to the conclusion that the main driver for the observed growth defects of GMSs is a naturally orchestrated catabolite uptake repression induced by the respective network perturbations. In detail, a disruption showing a phenotype implicates a decrease in the levels of one or more amino acid precursors, since evidently tight proteome coordination (You et al., 2013) hampers the enzyme allocation towards alternative precursor synthesis routes. According to the coarse-grained model of You et al. (2013) considering the general mechanism of ribosomal RNA transcription (Paul et al., 2004), low amino acid levels stall ribosomal protein synthesis via ppGpp. Moreover, low precursor levels, such as oxaloacetate or α-ketoglutarate, lead to an increased cAMP synthesis and elevated CRP-cAMP levels, which foster the expression of catabolic enzymes for alternative substrates and further reduce growth. Elevated cAMP concentrations, which were experimentally observed in multiple GMSs (McCloskey et al., 2018b), indicate increased CRP-cAMP activities and support this view on a gene deletion triggering the integral feedback of metabolic control.

**Figure 10.**
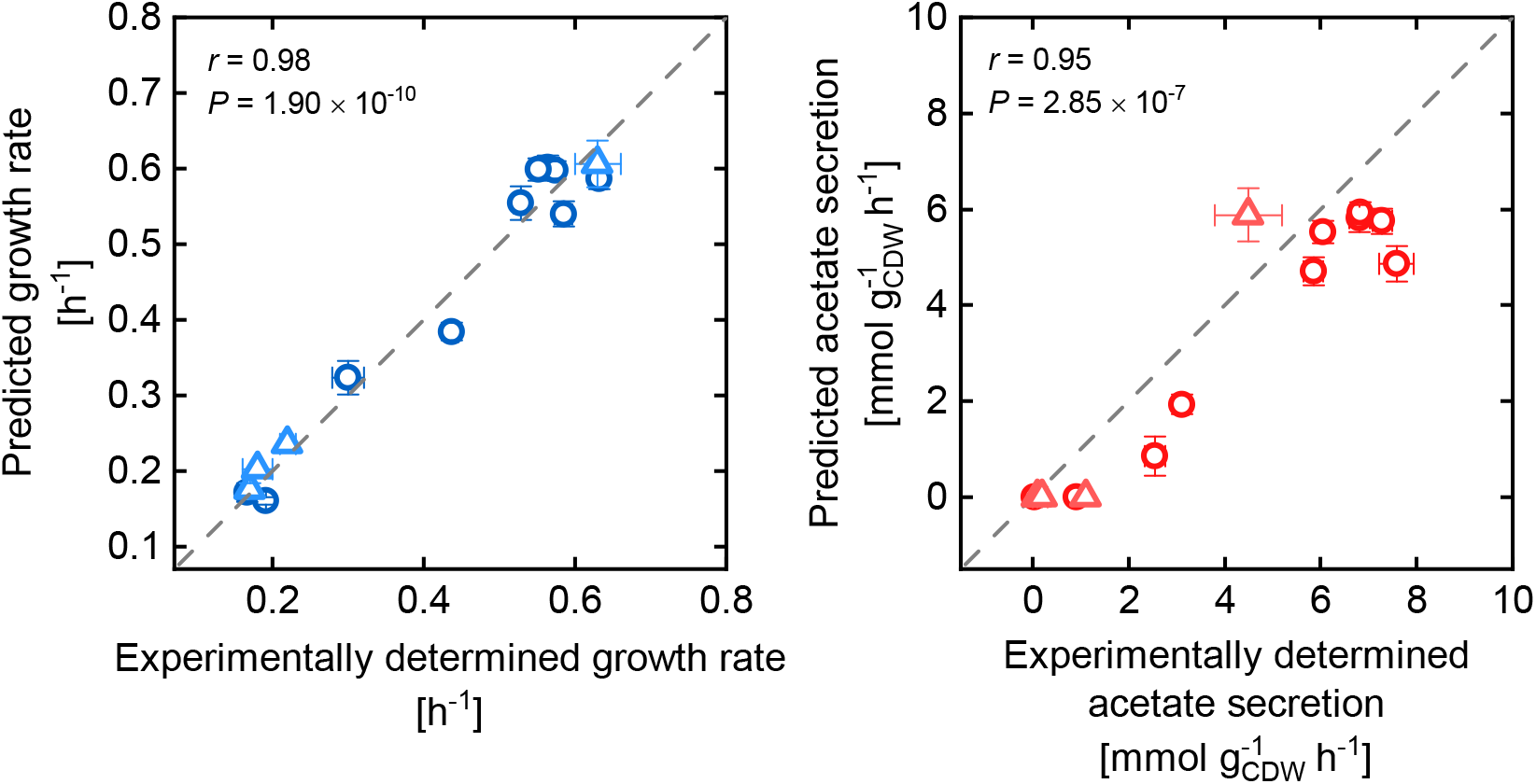
Comparison of predicted growth and acetate secretion rates of GMS with experimentally determined values taken from Long et al. (2016) and Fong et al. (2006). Predictions were made applying the PAM and constraining the glucose uptake rates to observed values.

The assumption that a regulatory substrate uptake inhibition shapes GMS’ phenotypes, raised the question if substrate uptake modes can be quantitatively predicted by the PAM. To tackle this question, we recalled the success of computational frameworks such as Minimization Of Metabolic Adjustment (MOMA) Segrè et al. (2002), Regulatory On/Off Minimization (ROOM) Shlomi et al. (2005) or RELAtive CHange (RELATCH) Kim and Reed (2012) in determining the metabolic impact of gene knockout strains. The common principle behind all three methods is the minimization of the metabolic response to genetic perturbations due to an unchanged regulatory system that forces the GMS’ flux distribution towards the original steady state. In the context of protein allocation, the minimal response principle can be translated as follows: Upon a network perturbation, a GMS establishes a substrate uptake rate so that *increases* in protein allocation towards single enzymatic reactions are minimal compared to genetically unperturbed strains. The cellular objective is to allow for maximum metabolic activity in the face of knockout-induced flux rerouting and a hampered reallocation of protein due to a strict (wildtype) regulatory regime.

To apply the minimal response principle on the PAM for mutant phenotype predictions, we implemented a strategy based on the analysis of growth optimal flux distributions at a range of substrate uptake rates (cf. Methods section for a detailed description). For each flux distribution, the difference in the enzyme synthesis rate Δ*N*_e_ between a reference, wildtype state at maximum growth and a mutant state is calculated for each enzyme within the PAM. The maximum substrate uptake rate of the GMS is determined as soon as Δ*N*_e_ meets a defined upper bound 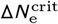 for one enzyme within that flux distribution. In doing so, we assign a certain flexibility to the overexpression capacity of single enzymes, which may be attributed to the utilization of unused enzymes. Moreover, if 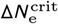 is met for an enzyme, flux rerouting to circumvent the saturated reaction or pathway is not possible.

Fig 11 shows the prediction results in comparison to experimentally determined phenotypic data (Fong et al., 2006; Long et al., 2016; Long and Antoniewicz, 2019b) for an 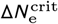 of 16 nmol 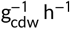 (cf. Methods section for the determination of 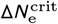). With one exception, there is a general agreement between predicted and experimentally determined phenotypes (Pearson correlation: *r* = 0.81, *P* = 7.76 × 10^-11^). By disregarding the *rpe* deletion mutant, the Pearson correlation improves to *r* = 0.97 with a *P* value of 1.5 × 10^-23^. For the omitted outlier, simulations show a near-wildtype behavior similar to the *gnd* GMS, whereas growth is significantly reduced in the experiments. This observed growth defect is peculiar, since knockouts of enzymes adjacent to ribulose-phosphate 3-epimerase do not show such a growth-limited phenotype. Moreover, the experimentally determined phenotype of an *rpe* GMS from Nakahigashi et al. (2009) was similar to the wildtype *E. coli* strain, thus obscures a plausible judgment of the simulated phenotype.

**FIGURE 11.**
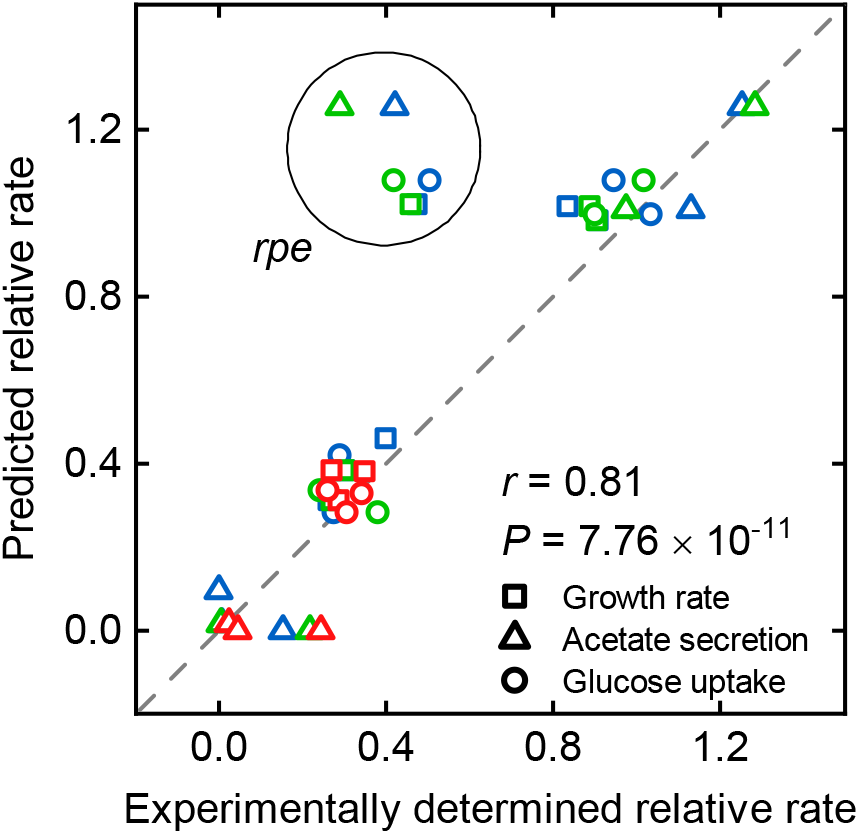
Comparison of predicted growth, glucose uptake, and acetate secretion rates of GMS with experimentally determined values taken from Long et al. (2016) (blue), Long and Antoniewicz (2019b) (green), and Fong et al. (2006) (red). All values were normalized with corresponding wildtype data. Predictions were made applying the PAM by constraining the excess enzyme sector according to a maximum glucose uptake rate of 9.82 mmol 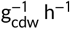. The maximum overexpression capacity of single enzymes 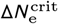 was set to 16 nmol 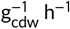.

The considerable agreement between experimental and simulated phenotypic data also translates to intracellular flux data, which we computed for single-gene deletions leading to a complete, demonstrable inactivation of the corresponding reaction (Long and Antoniewicz, 2019b). Predicted flux responses correlate well with predicted flux data from Long and Antoniewicz (2019b) (Fig 12), noticeable on Pearson’s correlation coefficients of *r* > 0.93 for all tested single deletions, except for the Δ*tpiA* mutant (*r* = 0.67). Concerning the Δ*tpiA* GMS, an experimentally observed rerouting of glycolytic flux through the methylglyoxal pathway towards pyruvate (McCloskey et al., 2018a; Long and Antoniewicz, 2019b) stands in contrast to the results of the PAM suggesting the activation of the ED pathway to surpass the blocked glycolysis. The simulated behavior can be traced back to an underestimated protein cost of the ED pathway. By decreasing the *k*_cat_ values of the two central ED pathway reaction steps phosphogluconate dehydratase and 2-dehydro-3-deoxyphosphogluconate aldolase to 2.5 % of the original values and following a re-evaluation of the maximum glucose uptake rate, glycolytic flux was diverted into the methylglyoxal pathway simultaneously leading to an activated acetate secretion (Appendix Fig S4). This selective adaption of turnover numbers, and thus of protein costs for the corresponding reactions, resulted in a significant convergence of simulated flux data towards measured values for the Δ*tpiA* and also the Δ*rpe* GMS (*r* of 0.90 and 0.99, respectively) (Appendix Fig S5). However, flux predictions for the Δ*gnd* mutant were deteriorated at the same time (*r* = 0.95). Here, flux was shifted from the ED pathway towards glycolysis, which contradicts experimental data. Nevertheless, the *k*_cat_ value adaption only slightly lessened the PAM’s overall predictive capabilities (Appendix Fig S5), thus generally points to the need for fine-tuning kinetic parameters to improve the flux split between pathways across different conditions.

**FIGURE 12.**
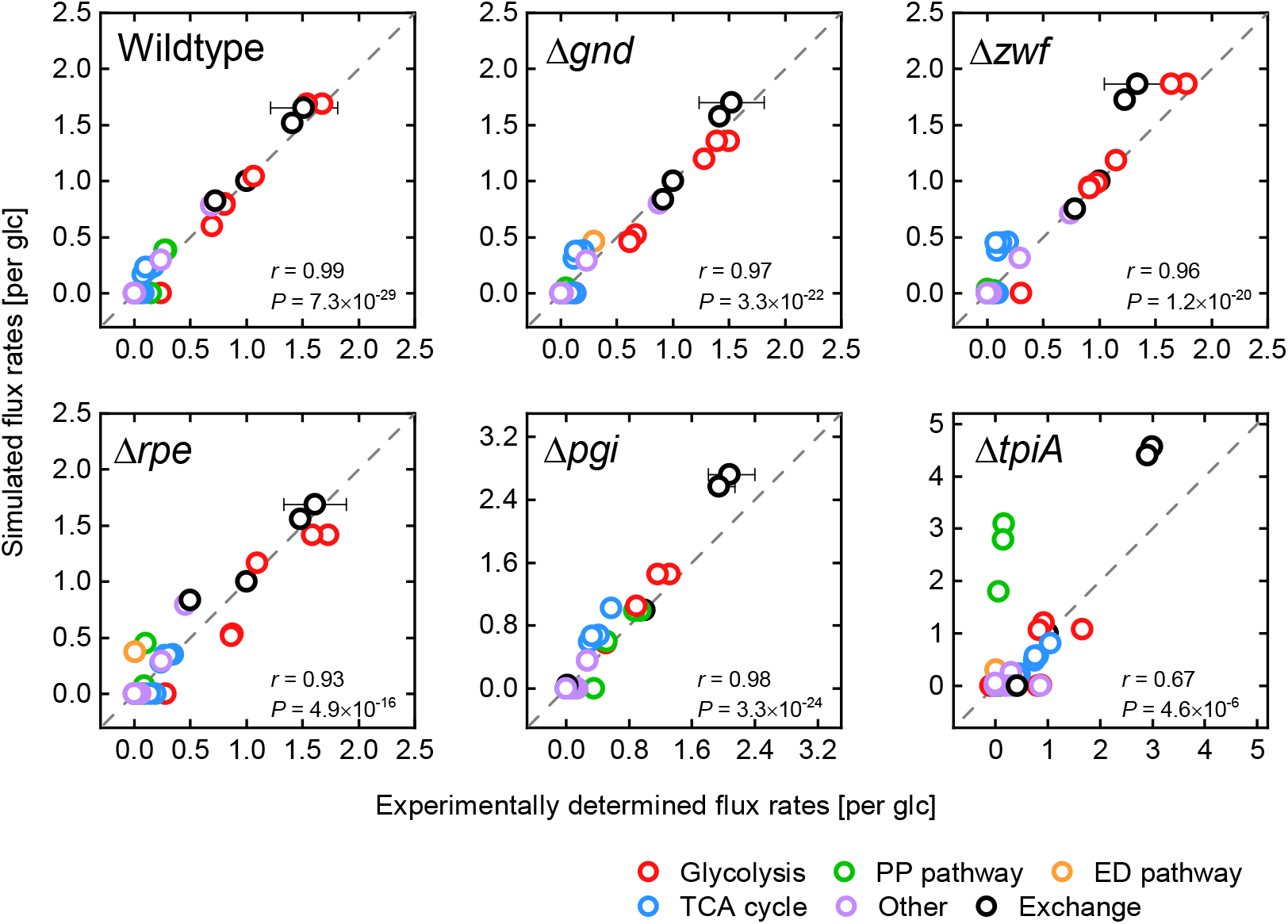
Comparison of predicted intracellular flux rates of GMSs with experimentally determined values taken from Long and Antoniewicz (2019b). All values were normalized with corresponding glucose uptake rates. Predictions were made applying the PAM by constraining the excess enzyme sector according to a maximum glucose uptake rate of 9.82 mmol 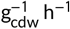. The maximum overexpression capacity of single enzymes 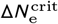 was set to 16 nmol 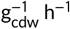.

## 3 DISCUSSION

Proteins are *the* major molecular class in cells and being the catalyst for global cellular functionalities, the importance of the mutual connection between microbial metabolic behaviors and protein allocation is well appreciated (Scott et al., 2010, 2014; Weiße et al., 2015; Mori et al., 2016; Noor et al., 2016; Erickson et al., 2017; Mori et al., 2019). Based on existent techniques for the consideration of protein allocation and enzymatic constraints in the well established constraint-based modeling area (Mori et al., 2016; Sánchez et al., 2017), we implemented a methodology to account for the total condition-dependent proteome in an *E. coli* GEM. Besides the integration of basic enzyme kinetics in the form of turnover numbers, the resulting PAM considers simple, linear relations between microbial growth and the ribosomal as well as the excess protein sector to describe 81 % of the total cellular protein cost.

The PAM’s accurate predictions confirmed the prominent role of protein allocation in shaping the microbial metabolism. Nevertheless, protein allocation itself appears to be regulated to exhaust fundamental biochemical limits. Maximum substrate uptake rates, and therefore also maximum growth rates, are dictated by an ultimately limited total synthesis rate of condition-dependent proteins. On a biochemical level, the apparent restriction of total protein copy numbers translates to a limited, cell-wide transcription capability. The causal link between transcription limitations and maximum proliferation rates is also indirectly suggested by genetic sequencing results of *E. coli* strains that have undergone extensive adaptive laboratory evolution (ALE) (LaCroix et al., 2015). The adapted strains, all exhibiting a fitness increase of up to 1.6 fold, showed mutations in the *rpoB* or *rpoC* gene leading to point mutations in the primary sequence of the *β*/*β*’ subunit of the RNA polymerase, which is part of the enzyme’s active center. Thus, the observed mutations globally affect transcription (LaCroix et al., 2015). Based on our study, we presume a corresponding reduction of transcriptional limitations allowing for an increased protein allocation towards the metabolically active enzyme and ribosomal protein sector. A detailed proteomics study is necessary to uncover changing protein allocation principles in growth-optimized microbial strains as well as to test this hypothesis with the PAM.

Recognizing transcriptional limitations as hard constraints for the microbial metabolism and by recalling that the presented PAM predictions represent growth optimal flux states, a cellular principle is deduced supporting previous findings (Mori et al., 2019): Particularly under substrate-limited conditions, *E. coli* regulates central cellular processes to maintain a Pareto-optimum between growth rate and the ability to flexibly adapt its metabolism to changing environmental conditions. The degree of flexibility is directly linked to the amount of allocated excess enzymes, which is hard-coded within the PAM. Interestingly, even when protein becomes a metabolically limiting factor, apparent from the onset of overflow metabolism, the cell maintains a strictly substrate uptake orientated allocation of protein to the excess enzyme sector. This preservation of flexibility could allow for an evolutionary advantage in the original environmental niche, however, may be a promising target for engineering *E. coli* or any microbe towards a high-performance cell factory.

Recently, systems metabolic engineering was emphasized as an integral part in the development and optimization of microbial cell factories (Wittmann and Lee, 2012; Maia et al., 2016; Saa et al., 2019; Choi et al., 2019; Sarkar and Maranas, 2019). Thus, we want to put the PAM forth to highlight the advantages of considering the allocation of the total condition-dependent proteome for constraint-based modeling techniques in favor of more accurate mutant strain predictions. We showed sound predictions of growth upon a range of overexpression levels of a non-enzymatic protein. The predictive capability for the overexpression of enzymes participating in metabolic reactions, e.g. in heterologously expressed pathways, still needs to be verified. However, the strict regulation of protein allocation, represented by the functional description of ribosomal and excess enzyme sectors in the PAM, appeared to shape metabolic responses and is insensitive to genetic interferences. We also confirmed a rather inflexible protein allocation behavior for gene deletion mutant strains. Coherent prediction results were obtained by allowing only small divergences from a wildtype state in the expression rates of single enzymes. Hence, we suggest, additionally to the CRP-cAMP mediated restrictions, a link to transcriptional limitations, similar to our aforementioned observation of maximum growth of wildtype strains. This hypothesis is supported by ALE experiments in which adaption of single-gene deletion mutants frequently generated mutations affecting the regulation of global and pathway-specific transcription (McCloskey et al., 2018b), possibly eliminating transcriptional hurdles and (partly) restoring growth rates.

In summary, we want to stress the importance of considering protein allocation constraints in GEMs for the systematic constraint-based reconstruction and analysis (COBRA), and the engineering of microbial metabolism without having to sacrifice computational speed or applicability of established COBRA methods (Heirendt et al., 2019). Beside the facilitated advantages, the limited availability and credibility of basic enzymatic kinetic data, particularly in the form of turnover numbers, still poses a major obstacle in providing PAMs for any microorganism. Therefore, we join the call for the establishment of a thorough kcatome as part of an accessible, genome-wide kinetome (Nilsson et al., 2017).

## 4 METHODS

### 4.1 Formulating and solving the protein allocation model

All flux solutions and corresponding phenotypes in this work represent growth optimal solutions of the following, classical FBA-problem including additional protein allocation and enzymatic constraints:

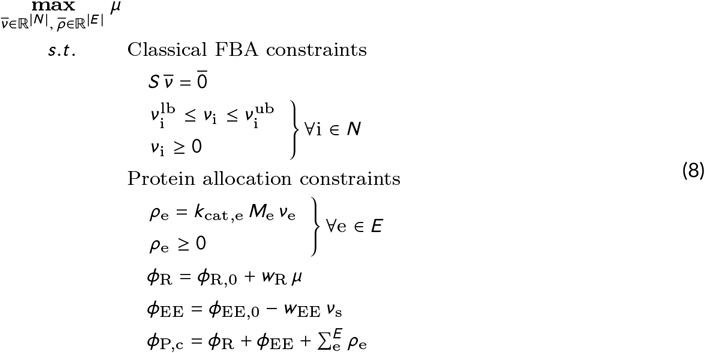

Here, *S* is the stoichiometric matrix of the original GEM and *ν*_i_ the flux variable of reaction i from the metabolic reaction pool *N*. Note that each reversible reaction in the original GEM has been split into irreversible forward and backward reactions to only allow for positive flux values, hence the lower and upper flux bounds 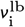 and 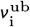 are equal or greater than zero. The additional protein allocation and enzymatic constraints comprise the mass concentrations *ρ*_e_ for each considered enzyme or enzyme complex, as well as the mass concentrations of the ribosomal and excess enzymes sector *φ*_R_ and *φ*_EE_, respectively. All protein allocation constraints in equation (8) were added to the stoichiometric matrix S of the basis GEM *i*ML1515 representing the *Escherichia coli* K-12 MG1655 strain (Monk et al., 2017).

Each reaction e from pool E, comprising all reactions that are linked to one or more genes via a gene-protein-reaction (GPR) relation, is assigned to exactly one protein with a unique turnover number *k*_cat,e_ and molar mass *M*_e_. In case a reaction e is catalyzed by an enzyme complex (multiple genes are connected via logical AND operators in the GPR relation) the molar mass *M*_e_ is the sum of molar masses of the participating gene products. If two or more enzymes are able to catalyze the same reaction independently from each other (multiple genes are connected via logical OR operators in the GPR relation), the isozymes are merged into one hypothetical protein with a molar mass equal to the mean of the molar masses of the merged isozymes. Molar masses of enzymes were calculated as the sum of molar masses of the amino acids constituting the respective primary sequences while taking into account the loss of one water molecule per peptide bond. The amino acid sequences were retrieved from the KEGG database (Kanehisa et al., 2017) by queries with the GEM-inherent, KEGG-specific gene identifiers. The *k*_cat_ values for all enzymatic reactions in the model were retrieved from BRENDA (Jeske et al., 2019) following a protocol of Sánchez et al. (2017). Since simulated maximum growth rates were unreasonably low when applying an initial *k*_cat_ set, a BRENDA-, SABIO-RK-(Wittig et al., 2012), and UniProt-based (Bateman, 2019) manual curation was done for those turnover numbers that showed the most pronounced effects on growth rates. The final set of curated *k*_cat_ values as well as molar masses of enzymes used throughout this study can be found in the Additional Files.

The ribosomal and excess enzymes sector *φ*_R_ and *φ*_EE_ are linearly related to growth rate μ and the substrate uptake rate *ν*_s_, respectively. As described in the Results chapter, the linear equations of both sectors (equation (8)) are parametrized according to thorough measurements and analyses of the total, condition-dependent proteome (O’Brien et al., 2016; Schmidt et al., 2016) and, except for the maximum substrate uptake rate *ν*_s_, are maintained for any simulations in this work. Finally, the total mass concentration of condition-dependent protein *φ*_P,c_ is kept constant and comprises the sum of the ribosomal sector *φ*_R_, excess enzymes sector *Φ*_EE_, and each considered enzyme *ρ*_e_. An overview of the applied parameters is given in Table 2.

**TABLE 2.**
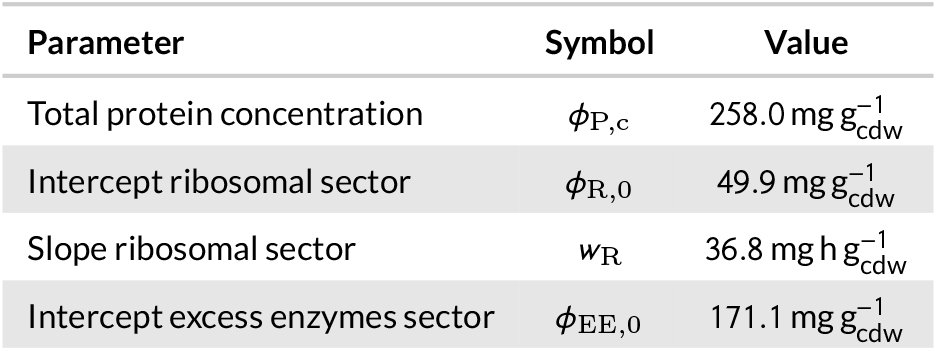
Universal PAM parameters applied for all simulations in this study.

| Parameter | Symbol | Value |
| --- | --- | --- |
| Total protein concentration | $\phi_{P,c}$ | 258.0 mg g<sub>cdw</sub><sup>-1</sup> |
| Intercept ribosomal sector | $\phi_{R,0}$ | 49.9 mg g<sub>cdw</sub><sup>-1</sup> |
| Slope ribosomal sector | $w_R$ | 36.8 mg h g<sub>cdw</sub><sup>-1</sup> |
| Intercept excess enzymes sector | $\phi_{EE,0}$ | 171.1 mg g<sub>cdw</sub><sup>-1</sup> |

#### 4.1.1 Determination of maximum substrate uptake rates

By applying the fully parametrized PAM, maximum uptake rates for single substrates were determined according to a maximally allowable total protein synthesis rate *N*_P_ (cf. Results chapter). Our hypothesis was that by parametrizing the slope w_EE_ of the excess enzymes sector so that maximum uptake rates for any substrate are met at a unique *N*_P_, the PAM should correctly predict phenotypes at substrate limited or measured conditions. Therefore, *N*_P_ values were calculated for substrates covered by the study of Gerosa et al. (2015) at a wide range of maximum substrate uptake rates based on growth-optimal flux distributions and while parametrizing the excess enzymes according to equation (4). Therefore, we generally assumed an absence of excess enzymes at maximum substrate uptake rates.

By using the respectively parametrized PAM, phenotypes are simulated for experimentally determined substrate uptake rates (Gerosa et al., 2015), if the assumed maximum uptake rates exceed the measured values. These simulated phenotypes, more specifically the growth rates, were then compared with the observed values by computing the absolute difference. For similar *N*_P_ values among the tested substrates, these absolute differences were summed up. At a *N*_P_ of 2.04 μmol 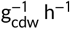 the sum of absolute differences between simulations and measurements was minimal (c.f. Fig 6) leading to maximum substrate uptake rates shown in Table 1.

### 4.2 Mutant strain simulations

#### 4.2.1 eGFP overexpression

Expression of eGFP was simulated by introducing an additional column to the stoichiometric matrix of the PAM representing a protein with mass of 2.8 × 10^4^ gmol^-1^. Thus, the expression strength of eGFP is controlled by the respective model variable describing the protein’s intracellular concentration. To identify the total protein concentration that represents the elevated protein availability in the *E. coli* TUNER strain used by Bienick et al. (2014), the summed absolute difference between measured and simulated relative growth rates at all experimentally tested eGFP expression strengths was computed for a range of total protein concentration *φ*_P,c_ values. The observed and simulated growth rates were normalized with wildtype growth rates. Student’s t-tests were applied to determine the *φ*_P,c_ yielding the best possible correlation between measurements and simulations.

#### 4.2.2 Phenotype determination of gene deletion mutants

The deletion of a gene was simulated by identifying all reactions connected to this gene via the model-inherent GPR and setting the respective upper bounds to zero. To predict maximum substrate uptake rates of GMSs, enzyme synthesis rates were calculated from growth optimal flux distributions for a wide range of substrate uptake rates. The synthesis rate *N*_e_ of an enzyme e was computed from flux distributions by

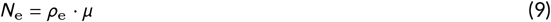

where *μ* and *ρ*_e_ are the growth rate and the molar concentration of enzyme e, both being optimization variables to the PAM.

For each tested substrate uptake rate the maximum difference in the computed enzyme synthesis rate 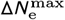 between the GMS and a reference state representing a wildtype strain grown under substrate-unlimited conditions was identified among all modeled enzymes. By scanning 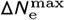 values from high to low substrate uptake rates, the maximum substrate uptake rate for a GMS is found as soon as 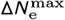 meets a critical value 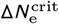 (also termed maximum overexpression capacity). Thus, the corresponding metabolic mode supports a maximally achievable growth rate under a constrained flexibility to change or reallocate protein among metabolic pathways and their single reactions. We assumed the level of restriction in flexibility 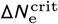 to be the same in any GMS according to phenotypic data of GMS from Long et al. (2016), Long and Antoniewicz (2019b), and Fong et al. (2006). We found that a 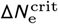 of 16.0 nmol 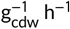 leads to a minimal sum of errors between predicted and observed growth, substrate uptake, and acetate secretion rates. Corresponding comparisons of experimentally determined and predicted phenotypes as well as flux distributions are shown in Fig 11 and Fig 12, respectively.

### 4.3 Implementation

All conducted simulations, model reconstructions, and data analyses were performed in MATLAB 2018a on a Windows 7 machine with 16 GB of RAM and an AMD FX-8350 Eight-Core (à 4.00 GHz) processor. COBRA toolbox functions (Heirendt et al., 2019) and the Gurobi Optimizer (8.0.0, Gurobi Optimization, Inc.) were utilized for processing and solving the metabolic models. All MATLAB functions necessary to handle and build a protein allocation model (PAM) from a COBRA format-based, stoichiometric reconstruction are provided on GitHub (https://github.com/Spherotob/_PAM_public).

## 5 ABBREVIATIONS

### List of abbreviations used

ALE: Adaptive laboratory evolution
CAFBA: Constraint allocation flux balance analysis
cAMP: Cyclic adenosine monophosphate
CDW: Cell dry weight
COBRA: Constraint-based reconstruction and analysis
COG: Clusters of orthologous groups
CRP: cAMP receptor protein
ED: Entner-Doudoroff
eGFP: Enhanced green fluorescent protein
EMP: Embden-Meyerhof-Parnas
GEM: Genome-scale model
GMS: Gene-deletion mutant strains
GPR: Gene-protein-reaction
ME: Macromolecular expression
PAM: Protein allocation model
PEP: Phosphoenolpyruvate
PP: Pentose phosphate
ppGpp: guanosine 5-diphosphate 3-diphosphate
PTS: Phosphotransferase system
TCA: Tricarboxylic acid

## 6 ACKNOWLEDGMENTS

TBAwas partially funded by the Excellence Initiative of the German federal and state governments (Grant-ID: PFLS015). BEE acknowledges partial funding through the CSIRO-UQ Synthetic Biology Alliance. The laboratory of Lars M. Blank was partially funded by the Deutsche Forschungsgemeinschaft (DFG, German Research Foundation) under Germany’s Excellence Strategy within the Clusters of Excellence 236 “TMFB” and 2186 "The Fuel Science Center”. The funders had no role in the design of the study and collection, analysis, and interpretation of data and in writing the manuscript.

## 7 AUTHOR CONTRIBUTION

TBA and BEE conceived and designed the study. TBA conducted the computational implementations, performed all simulations, and evaluated the simulation results. TBA and BEE wrote the manuscript. LMB supervised the study and edited the manuscript. All authors read and approved the final manuscript.

## 8 CONFLICT OF INTEREST

The authors declare that they have no conflict of interest.

## 9 DATA AVAILABILITY

The basic Matlab implementations used in this study can be found on GitHub: https://github.com/Spherotob/PAM_public. All relevant data used to draw the Figures in this manuscript is presented in Additional File 1. Details of the metabolic model and the set of curated *k*_cat_ values is listed in Additional File 2. Supplementary figures can be found in Additional File 3.

## LIST OF APPENDIX FIGURES

S1 Comparison between experimentally determined and predicted maximum substrate uptake rates, as well as the corresponding growth and acetate secretion rates using several substrates as sole carbon sources.

S2 Comparison of predicted growth and acetate secretion rates of GMS with experimentally determined values taken.

S3 Comparison of phenotypic predictions between wildtype and gene-deletion mutant model.

S4 Comparison of predicted growth, glucose uptake, and acetate secretion rates of GMS with experimentally determined values at decreased kcat values of Entner-Doudoroff pathway reactions.

S5 Comparison of predicted intracellular flux rates of gene-deletion mutant strains with experimentally determined values at decreased kcat values of Entner-Doudoroff pathway reactions.

